# UVA irradiation promotes ROS-mediated formation of the common deletion in mitochondrial DNA

**DOI:** 10.1101/2025.07.09.663484

**Authors:** Gabriele A. Fontana, Navnit K. Singh, Nadezhda Rotankova, Antonia Eichelberg, Michela Di Filippo, Michael Mc Arthur, Susanne Heldmaier, Franziska Wandrey, Hans-Dietmar Beer, Shana J. Sturla, Hailey L. Gahlon

## Abstract

Ultraviolet (UV) radiation from the sun causes adverse skin changes such as premature aging. UVA radiation is the primary factor for photoaging due to its deep penetration into the dermis, and UV-induced mitochondrial DNA (mtDNA) alterations, including deletions, contribute to photoaging and cellular dysfunction. The most frequent mtDNA rearrangement is the common deletion (CD), characterized by the loss of nearly one-third of the genome, 4,977 base pairs. UV radiation exposure leads to the formation of the CD, however, a distinct characterization of UV-induced CD and the underlying molecular mechanisms driving its initiation remains unexplored. In this study, we showed that increasing doses of UV radiation led to an increase in the CD in human skin fibroblasts. We found that UVA induce the formation of the CD by increasing the cellular reactive oxygen species (ROS) and oxidized bases content in the mtDNA. Preconditioning cells with antioxidants prevented the accumulation of the UVA-induced CD, suggesting that this mutational mechanism is ROS-dependent. In stark contrast, UVB did not alter cellular ROS levels but increased the formation of cyclobutane pyrimidine dimers (CPD), leading to CD generation though a ROS-independent mechanism. We corroborated our findings by using a 3D human full-thickness skin equivalent model, where we detected UVA-dependent CD formation in both the epidermal and dermal layers of the skin. By analyzing bulk RNA from UVA-exposed human skin fibroblasts by RNA-Seq, we found that UVA led to the upregulation of genes encoding mitochondrial DNA replication proteins and to the downregulation of genes involved genes encoding mtDNA repair factors. Taken together, our findings provide insight into how UVA and UVB differ in their detrimental effects on mtDNA, with UVA impacting mtDNA maintenance and transcription via a ROS-dependent mechanism. Our findings also established the mtDNA CD as a novel potential biomarker for monitoring UVA-induced oxidative stress and photoaging in skin cells in vitro and in vivo.

## Introduction

Solar radiation exposure can damage skin, causing, skin cancer (1–3), as well as premature skin aging (4–6). UVA radiation (320-400 nm) is considered the primary cause of photoaging because it penetrates through the epidermis to the dermis, a connective tissue with collagen-producing fibroblasts required for the skin’s structural integrity (7). UVB (280-320nm) has a lower penetration depth, reaching the basal layer of the epidermis, but also has direct and indirect effects on both dermal and epidermal layers of skin (8). UVA exposure generates reactive oxygen species (ROS), such as hydrogen peroxide, hydroxyl radicals, and singlet oxygen, and prolonged exposure can overwhelm the body’s antioxidant defenses and induce oxidative stress, leading to cellular damage. UVA forms oxidative lesions, particularly 8-oxo-7,8-dihydro-2’-deoxyguanosine (8-oxo-dG), whereas UVB forms cis-syn-cyclobutane dimers and (6–4) pyrimidine photoproducts (9), making both potent mutagens that damage genomic DNA (10,11). However, the molecular pathways through which UV-induced ROS contribute to photoaging remain only partially elucidated. (12,13).

Mitochondria are organelles with a primary function of generating energy for the cell in the form of ATP via the electron transport chain, making them the site of the highest ROS turnover in cells (14). In humans, each mitochondrion contains several copies of a circular DNA molecule named mitochondrial DNA (mtDNA) of 16,569 base pairs, which encodes for 37 genes including 13 oxidative phosphorylation proteins, 22 tRNAs, 2 rRNAs, and the Humanin micropeptide. Mutations in mtDNA, including point mutations and small- and large-scale deletions, can disrupt mitochondrial physiology and lead to cellular dysfunction (15). Mitochondrial DNA (mtDNA) exists in a state of heteroplasmy, in which wild-type (non-mutated or non-deleted) and mutated or deleted mtDNA molecules coexist within the same cell (16). UV-induced mtDNA mutations have been found in photoaged human skin cells (17–19) and large-scale deletions of the mitochondrial genome are implicated in UV radiation-induced photoaging of the skin (19–24). Mitochondria have been found to play a critical role in skin aging (17,25,26) and are recognized as one of the hallmarks of aging (27), making them a viable target for anti-aging interventions (28,29).

The common deletion (CD) is the most prevalent large scale mitochondrial deletion involving the removal of almost a third (4,977 base pairs) of the mitochondrial genome. The mtDNA region deleted by the CD is flanked by two 13-base-pair repeats, with one repeat being retained in the deletion. This deletion was first reported in 1989 in a patient with Kearns-Sayre syndrome (KSS) (30). UV light appears to promote formation of the mtDNA CD (24), and higher levels of CD were observed to form after UV-exposure of skin and are associated with photoaging (22,31-34). We previously suggested the CD as a candidate biomarker for cancer-associated fibroblasts (35). To explore the potential of the CD as a biomarker for UVA-induced photoaging, we require molecular characterization of UV-induced CD formation and evaluation of its ability to respond to possible interventions, such as antioxidants, in the case of ROS-dependent formation.

Several theories have been proposed regarding details of how the CD in mtDNA forms, including the replication-slippage model (30), the copy-choice recombination model (36), and DNA repair-associated deletion formation (37,38). The replication-slippage model and the copy-choice recombination model depend on replication of mtDNA, which consists of a heavy strand (H-strand) and a light strand (L-strand) that are replicated from their respective origins of replication (39). The replication-slippage model proposes that fork stalling during H-strand synthesis promotes mis-annealing of single-stranded mtDNA regions. The 13-base pair 3’ repeat of the H-strand mis-anneals with the downstream 5’ repeat, forming a loop that can be extruded from the mtDNA leading to the formation of the mtDNA CD. Interestingly, a modified replication-slippage model suggests that singlet oxygen generated, for example by UVA exposure, induces base damage in guanine-rich regions, leading to strand breaks necessary for the degradation of the looped-out DNA (29). The formation of the CD through the copy-choice recombination model involves the synthesis of the mtDNA L-strand. It is proposed that mtDNA deletions can be formed during the repair of double-strand breaks through mechanisms such as homologous recombination or nonhomologous end-joining, with the latter potentially contributing to deletions lacking direct repeats (37,38,40). Given the overlap between replication and repair pathways in the mitochondria, it is also possible to envision a mechanism that relies on replication-repair crosstalk (15,41). While these mechanisms could explain how the mtDNA CD is formed, involving oxidative stress (42,43) and mtDNA replication fork stalling (15), data linking how UV radiation initiates the formation of the CD are still lacking.

In this study, we characterized the degree of formation of the mtDNA CD in human skin fibroblasts following UVA or UVB exposures. To probe the role of oxidative stress in the initiation of CD formation, we simultaneously monitored changes in ROS and base damage in mtDNA. Furthermore, we supplemented fibroblasts with antioxidants in combination with UVA exposure to evaluate its impact on cellular ROS and how it related to the formation of the CD. By assessing changes in gene expression in UVA exposed fibroblasts, we identified clusters of genes that are up- and downregulated in response to UVA irradiation. This study reveals how UVA and UVB differentially induce CD formation and highlights its correlation with oxidative stress. Additionally, it shows that UVA specifically shapes transcriptional responses, providing new insights into the mitochondrial contribution to photoaging.

## Material and methods

### 2D fibroblast cell culture

BJ-5ta human foreskin fibroblasts (ATCC, CRL-4001™) were maintained in a 4:1 mixture of Dulbecco’s DMEM high glucose (DMEM; ThermoFischer Scientific) and Medium 199 (ThermoFischer Scientific), and supplemented with 10 % fetal bovine serum (FBS; ThermoFisher Scientific) and 0.01 % mg/ml Hygromycin B from Streptomyces hygroscopicus (Sigma-Aldrich). KSS human skin fibroblasts from a donor with Kearns-Sayre Syndrome were enucleated and fused to 143B osteosarcoma cells, and were kindly provided by Professor Ivan Tarassov (University of Strasbourg) (43). KSS fibroblasts were cultured in DMEM (ThermoFischer Scientific) supplemented with 10 % FBS (ThermoFischer Scientific) and 1 % Penicillin-Streptomycin (ThermoFischer Scientific). Both cell lines were cultured at 37 °C in 5 % CO_2_. Cells were routinely passaged every 2-3 days. For passaging, cells were washed once with Dulbecco’s phosphate-buffered saline (DPBS, ThermoFischer Scientific), then detached with Trypsin-EDTA (0.25%), phenol red (ThermoFisher Scientific). Trypsin was neutralized with the corresponding medium and cells were passaged in a 1:XYX proportion in a new Petri dish.

### Generation of human skin equivalents

Skin equivalents were generated per the protocol described by Berning, Prätzel-Wunder et al. 2015 (44). In summary, for the dermis-like compartment, human dermal fibroblasts (HDF) were seeded onto 12-well translucent ThinCerts™ (high-density 0.4 μm pores, 11.31 cm^2^ culture area, VMR) at a density of 5×10^5^ cells per well on days 1, 3, and 5 and placed in deep-well plates (VMR). The HDF were incubated at 37 °C, 5 % CO_2_ and 20 % O_2_ over a total culture time of 4 weeks with a resulting input of HDFs of 1.5×10^6^ cells per transwell insert. HDFs were cultured in 3:1 DMEM/Ham’s F12 (FAD) containing 10 % FBS (Sigma) and 1 % Antibiotic-Antimycotic (100x) (Thermo Fisher Scientific), supplemented with 200 μg/mL 2-phospho-l-ascorbic acid (Sigma) and additional recombinant human proteins: 1 ng/mL transforming growth factor β-1 (TGFβ-1; Thermo Fisher Scientific), 2.5 ng/mL epidermal growth factor (EGF; Thermo Fisher Scientific), 5 ng/mL basic fibroblast growth factor (bFGF; Peprotech), and 5 μg/mL insulin (Sigma). Medium was changed every other day, for 4 weeks. For the epidermis-like compartment, 2.5 × 10^5^ human primary keratinocytes (HPK) were seeded per dermal equivalent in FAD medium containing 10 % FBS and 1 % antibiotics/antimycotics, supplemented with 200 μg/mL 2-phospho-l-ascorbic acid-trisodium salt, 0.4 μg/mL hydrocortisone (Sigma), and 1×10−10 M cholera toxin (Sigma). After 3 days of submersed growth, the epithelial cocultures were air-lifted and the medium changed every other day, for 2 weeks.

### UVA and UVB irradiation

Cells were grown to 70-80% confluency in 15 cm Petri dishes. Prior to irradiation, the culturing medium was changed to serum-free DMEM (ThermoFischer Scientific) without antibiotics. Cells were irradiated in a UV irradiation chamber (BS-02 Opsytec Dr. Grödel) according to the schemes shown in the corresponding figures (Figures 1a, 3a, 5a, 5f). For UVA, dishes were covered with a soda-lime glass (150 × 25 mm, Faust Laborbedarf) to filter the remaining minor fraction of UVB emitted by the irradiation system. A dosimeter was place internally to the irradiation chamber and was used to monitor the UVA dose. For UVB exposure, irradiation for 3 min 20 s with four 5 mV UVB lamps (Opsytec) resulted in an irradiation exposure level of 0.5 J/cm^2^. UVB exposures were performed without a cover on the dishes. For experiments involving antioxidants, reduced L-glutathione (Sigma-Aldrich) dissolved in H_2_O (final concentration 50 µM) or coenzyme Q10 (CoQ10, Sigma-Aldrich) in dimethylformamide (DMF; final concentration 500 µM) were added to the serum-free medium. Serum-free medium was replaced with culturing medium after irradiation. During irradiation, all control dishes were kept at room temperature within a laminar flow hood.

**Figure 1.**
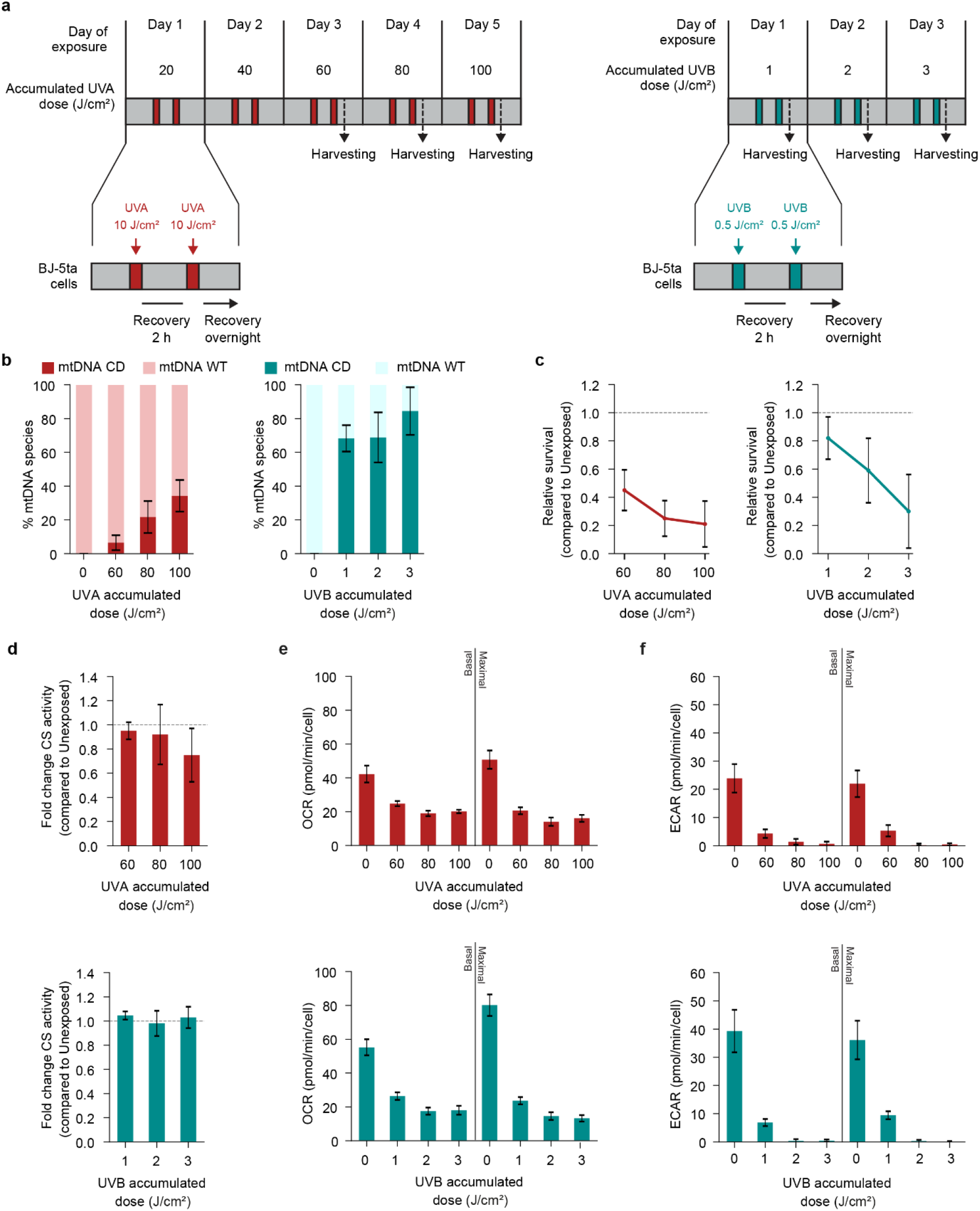
UV light-dependent induction of the mitochondrial DNA common deletion (mtDNA CD) in human fibroblasts. **(a)** Schematic representations of the UVA (left) and UVB (right) exposures performed on the BJ-5ta human fibroblast cell line. Cells in medium devoid of FBS and antibiotics were irradiated with UV twice per day with the indicated doses of UVA (red bars) or UVB (cyan bars). During recovery intervals (grey bars) cells were placed in an incubator with medium containing FBS and antibiotics. Harvesting was performed 30 minutes after the last indicated irradiation. **(b)** Relative quantification of wild-type mtDNA (mtDNA WT, light bars) and mtDNA harboring the common deletion (mtDNA CD, dark bars) using qPCR in Unexposed and UVA-(left) or UVB-exposed (right) cells. **(c)** Relative survival of BJ-5ta cells irradiated with indicated doses of UVA (left) or UVB (right) compared to Unexposed cells (grey dotted line). (**d)** Citrate synthase activity in UVA-(top) and UVB-exposed (bottom) BJ-5ta cells. The citrate synthase activity in UVA and UVB exposed cells is expressed as fold-change per well to Unexposed control cells (grey dotted line). **(e)** Oxygen consumption rates (OCR) at basal and stimulated (maximal) ETC activity in UVA (top) and UVB (bottom) exposed cells. **(f)** Extracellular acidification rates (ECAR) at basal and maximal ETC activity in UVA-(top) and UVB-(bottom) exposed cells. For **(b-d)** n = 3 biological replicates were used and **(c-d)** and for **(e-f)** n = 7 biological replicates. Error bars represent standard deviation.

### Human skin equivalent staining

Skin equivalent samples were fixed overnight in formalin 4 %, dehydrated automatically and embedded in paraffin (Leica EG 1150, Leica microsystem). Paraffin blocks were cut into 5 μm-thick sections and stained with heamatoxylin and eosin (H&E) using the HistoCore SPECTRA ST according to its standard protocols or analyzed by immunohistochemistry or immunofluorescence.

### Immunohistochemistry

For specific staining of paraffin sections, tissue sections were deparaffinized in xylene and rehydrated by incubating in decreasing concentrations of ethanol (100 % - 96 % - 80 % - 70 % and 50 %) for 2 minutes, followed by water for 2 minutes. The activity of endogenous peroxidase was blocked by 10 minutes of incubation in 3 % peroxidase-blocking solution (hydrogen peroxide 30 % in ddH2O; Merck). For antigen retrieval, slides were heated in a steamer for 30 min in antigen retrieval solution, sodium citrate buffer, pH 6.0 (10 mmol/L sodium citrate, 0.05 % Tween20). Sections were then blocked for 1 h at room temperature with 5 % bovine serum albumin in PBST (0.05% Tween 20 in PBS, blocking solution) and incubated overnight at 4 °C with the primary antibody (Ki67 Abcam ab833, rabbit, 1:100) diluted in the blocking solution. Biotinylated secondary antibody (Anti-Rabbit IgG BIOT 4010-08 (Southern Biotech)) was applied for 1 h at room temperature. Staining was visualized with the AEC substrate (Dako); nuclei were stained using Mayer’s hematoxylin (Kantonsapotheke, Zürich, Switzerland). Slides were mounted with Faramount aqueous mounting medium (Dako) and scanned using an Aperio ScanScope (Leica Biosystems) or with the PhenoImager Vectra Polaris (Akoya Biosciences).

### mtDNA and RNA extraction

Cells were harvested by scraping and pelleted by centrifugation (3 min, 4000 rcf, at room temperature). Cell pellets were either processed immediately or stored at −20 °C. RNA was isolated from cell pellets using the RNeasy Mini Kit (Qiagen) following manufacturer’s instructions. The RNA preparation included an on-column RNase-Free DNase (Qiagen) treatment prior to elution. mtDNA was isolated from cell pellets as previously described (45). Concentrations of purified RNA and mtDNA were assessed using a NanoDrop (ThermoFisher Scientific) spectrophotometer. For mtDNA extraction from HSE, epidermal and dermal layers were separated, cells were lysed in Buffer P1 supplememted with RNAse (Qiagen) using TisseLyser (Qiagen) for 3 min at 30 Hz. The lysates were then processed similarly to the cell pellets, as described above

### Quantitative real-time PCR (qPCR) for quantitation of the CD

Purified and RNA-free mtDNA was quantified for the CD by qPCR as described previously (35,45). In brief, for each sample 5-10 ng mtDNA, 0.4 μM forward primer, 0.4 μM reverse primer, 6 μL GoTaq 2x qPCR Master Mix (Promega), and nuclease-free water were prepared to a final volume of 12 μL per sample. Three technical replicates were analyzed for each mtDNA and primer combination. All primers used in this study are listed in Table S1 and were obtained from Eurogentec. All qPCRs were conducted on the Rotor-Gene 6000 (Corbett Research) device, following a standard qPCR protocol. To obtain the relative content of mtDNA CD and undeleted species, qPCR data was normalized using the delta-delta Ct method using the amplification of the total mtDNA content in each sample as normalizer. Corresponding qPCR values for mtDNA CD and undeleted species were expressed as relative fractions to the corresponding total mtDNA. Genomic DNA contamination was routinely assessed by qPCR on a genomic region of the *ACTB* gene, which resulted in low or no signal.

### Quantitative real-time PCR (qPCR) for the analysis of gene expression levels

Purified RNA expression was quantified by qPCR as described previously (35). In summary, the High-Capacity cDNA Reverse Transcription Kit (ThermoFisher Scientific) was used to reverse transcribe 1-2 μg purified RNA into cDNA according to the manufacturer’s instructions. For each sample 4-6 ng cDNA, 6 µL HOT FIREPol EvaGreen qPCR Mix (Solis Biodyne), 0.4 μM forward primer, 0.4 μM reverse primer, and nuclease-free water were prepared to a total volume of 12 μL. All primers used in this study are listed in Table S1 and were obtained from Eurogentec, Belgium. All qPCRs were conducted on the Rotor-Gene 6000 (Corbett Research) device, using the following protocol for each cycle: 95°C for 30 seconds, 60 °C for 1 minute, 40-50 cycles, with fluorescence acquisition at the end of the last amplification step. Expression levels for each biological replicate were assessed using the delta-delta Ct method. For nuclear genome-encoded target genes, amplification of *GAPDH* or *ACTB* housekeeping genes were used as normalizer. For mtDNA-encoded target genes, the amplification of an untranslated region of the polycistronic mtDNA-encoded RNAs was used as normalizer.

### RNA-sequencing

RNA sequencing was performed by the Functional Genomics Center Zürich. RNA was extracted using the Quiagen RNeasy Mini Kit following manufacturer’s protocol. Extracted RNA was prepared for sequencing using the Illumina TruSeq Stranded mRNA Library Prep assay following manufacturer’s protocol. Sequencing was performed on the Illumina NovaSeq 6000 using the S1 Reagent Kit v1.5 (100 cycles) as per manufacturer’s protocol. Demultiplexing was performed using the Illumina bcl2fastq Conversion Software. Individual library size ranged from 20 million to 26 million reads.

### Measurement of reactive oxygen species (ROS)

1.5 × 10^4^ cells were seeded in 96-well-plates and assayed at a confluency of 70-80 %. Culturing medium was removed from each well and replaced with 200 μL of 50 μM 2′,7′-Dichlorofluorescin diacetate (DCF, Sigma Aldrich) solution in PBS (1x). Plates were incubated for 30 min (37 °C, 5 % CO_2_), the solution was removed, and 30 μL Trypsin (ThermoFisher Scientific) was added. Following 3-5 minutes incubation for detachment of the cells, 120 μL PBS (1x) were added and fluorescence was acquired using a CytoFLEX flow cytometer (Beckman Coulter). Data were normalized internally by gating. To determine the ROS fold change, the intensity of the cell-free blank control was first subtracted from each condition. Fold-change was determined from the ratio of intracellular ROS levels in UV-irradiated versus unexposed cells. Each condition was assayed in three biological replicates, and each sample was tested in three technical replicates.

### Citrate synthase assay

1.5×10^4^ cells were seeded in 96 well-plates and assayed at a confluency of ∼70-80 %. The activity of the citrate synthase enzyme, a proxy for cellular mitochondria content, was measured using the Citrate Synthase Activity Assay Kit (Abcam) following manufacturer’s instructions. The colorimetric readings were acquired using a microplate Tecan instrument (Tecan, Infinite 200 PRO model) and normalized to a cell-free blank and to the amounts of harvested cells for each sample. Each condition was assayed in three biological replicates, and each sample was tested in three technical replicates.

### Seahorse assay

Basal and UV-induced changes in oxygen consumption rate (OCR) and extracellular acidification rate (ECAR) were measured using a Seahorse XF96 Extracellular Flux Analyzer (Agilent Technologies). Assays were performed in Seahorse XF Assay Medium (pH 7.4) supplemented with glucose (10mM) and sodium pyruvate (1mM) and L-glutamine (2mM). Four measurement cycles were performed for each injection with mix, wait and measure durations set to 3, 2 and 3 minutes. A basal measurement was performed followed by oligomycin (1 μM), 4-(trifluoromethoxy)phenylhydrazone (FCCP, 1.5 μM) and rotenone/antimycin A (2 μM) injections. All OCR and ECAR values were normalized to cell count as measured by nuclear staining with Hoechst 33342 (Thermo Fisher) followed by automated cell counting using a Cytation 5 Cell Imaging Reader (Biotek) and Gen5 software (Biotek).

### Dot Blot

Dot blots were performed on mtDNA extracted from UV-irradiated and unexposed cells as a qualitative indication of the presence of 8-oxo-dG and CPD in the mtDNA. All reagents used in this procedure, from the extraction to the solutions used until the incubation with the primary antibody, were supplemented with 350 μM 8-hydroxyquinoline (Sigma, stock dissolved in 1:1 EtOH;H_2_O), an antioxidant used to prevent artifactual formation of oxidized mtDNA bases. Unless stated otherwise, all steps of this procedure were performed at room temperature. 2 μl of isolated mtDNA, corresponding to 20 ng of mtDNA, were spotted on pre-cut nitrocellulose membranes (Thermofisher scientific) and dried for 2 h at 60 °C. The samples were then immerged in 5 mL of blocking buffer, composed of 0.5 % BSA (Thermofisher scientific) in TBS-T (20 mM tris-HCl, 150 mM NaCl pH 7.5, 0.05 % Tween20) (Thermofisher scientific), and incubated for 1 h on a shaker. The blocking buffer was then removed, and the membranes were covered in a solution containing the respective primary antibodies (anti-ds DNA (ab27156, Abcam), anti-CPD (Clone TDM-2, Cosmo Bio, USA), anti-8-OHdG Antibody (15A3) (sc-66036, Santa Cruz Biotechnology). The samples were then kept overnight on a shaker at 4°C. From this point on, the solutions used were no longer supplemented with 8-hydroxyquinoline. The membranes were washed 3 times with TBS-T for 10 min, and then incubated for 1 h with the secondary antibody (Abcam ab6789, anti-mouse HRP-conjugated raised in goat, 1:2000) in TBS-T containing 0.5 % BSA. Then, the membrane was placed on a plastic foil and covered with mixed ECL reagent (Promega) for 5 minutes. HRP activity was then recorded using a ChemiDoc Imaging System (Bio-Rad). Uncropped membrane scans are provided in the corresponding supplementary figures.

### Primer extension experiments

To evaluate if the Pol G stalls at 8-oxoG, we performed a primer extension assay. For that, first 1.5 µL Pol G (1 µM), 5 µL 10x Pol G buffer (250 mM Tris (pH 7.5), 400 µg/mL, 100 mM MgCl_2_) and 5 µL DNA (1 µM) were pre-incubated at 25 °C for 5 minutes. To initiate the reaction 37.5 µL H_2_O and 1 µL dNTPs (5 mM) were added.

### Statistical analysis

All values are expressed as mean ± standard deviation (SD). Information on the number of biological replicates is given in the figure legends. Differences in mean values between groups were analyzed using GraphPad Prism 10 (GraphPad Software, USA), and the statistical analyses performed are indicated in the figure legends. In the case of significant differences, the P values obtained are shown in the panels, with P < 0.05 considered statistically significant.

RNA sequencing analysis was performed using the SUSHI framework (Hatakeyama et al., 2016), which encompassed the following steps: Read quality was inspected using FastQC, and sequencing adaptors removed using fastp; Alignment of the RNA-Seq reads using the STAR aligner (Dobin et al., 2013) and with the GENCODE mouse genome build GRCh38.p13 (Release 32) as the reference (Frankish et al., 2021); the counting of gene-level expression values using the ‘featureCounts’ function of the R package Rsubread (Liao et al., 2013); differential expression using the generalised linear model as implemented by the EdgeR Bioconductor R package, and; Gene Ontology (GO) term pathway analysis using both the hypergeometric over-representation test via the ‘enricher’ function, and gene-set enrichment analysis via the ‘GSEA’ function, of the clusterProfiler Bioconductor R package (Yu et al., 2012). All R functions were executed on R version 4.0

## RESULTS

### UVA and UVB induce CD formation in human skin fibroblasts

First, we evaluated how CD levels changed in response to UVA vs UVB exposure in cultured human skin fibroblasts. By a qPCR assay that we previously developed, we simultaneously measured CD, wild type (WT), and total mtDNA (45). Under exposure regimes (Figure 1a) consisting of sequential short exposures and longer recovery periods, with maximum irradiation of 100 J/cm^2^ UVA and 3 J/cm^2^ UVB, which were reached on day 5 for UVA or day 3 for UVB. For both UVA and UVB exposures, we observed a dose-dependent increase in the CD (Figure 1b), with UVA-induced CD levels ranging from 6 to 35 % for 60-100 J/cm^2^ (Figure 1b, left). As a control, we also measured levels of the UV responsive genes *MMP1* and *COL1A1*, which are reported to be upregulated and downregulated, respectively, following UVA exposure of fibroblasts (46), and by RT-qPCR analysis, we could confirm the expected differential regulation expression in gene expression (Figure S1). Thus, we found that CD levels following UVB exposure were higher, 70-80 % for 1-3 J/cm^2^, as compared to following UVA exposure (Figure 1b, right), suggesting different initiation processes or kinetics. To irradiate cells with a dose of UVA and UVB that would affect comparable rates of survival, we evaluated changes in cell survival following UVA or UVB exposure (Figure 1c). For UVA, a reduction in cell survival was observable at 60, 80 and 100 J/cm^2^, corresponding to day 3 to 5 of exposure (Figure 1c, left). A more notable and rapid decrease was observed for UVB in comparison to UVA between day 1 and 3 of UV irradiation (Figure 1c, right). However, maximum irradiation levels of 100 J/cm^2^ for UVA and 3 J/cm^2^ for UVB resulted in a mean cell survival of ∼30 % for both irradiation regimens. From this dose-response course, we defined the maximum doses of UVA and UVB that affect cell survival to a similar degree.

To test whether the accumulation of the mtDNA CD induced by UVA and UVB are accompanied by changes in mitochondrial function, we evaluated several parameters. First, we monitored cellular mitochondrial content using citrate synthase assay. Citrate synthase is an essential mitochondrial enzyme in the tricarboxylic acid cycle (TCA) cycle that catalyzes the formation of Citrate from Acetyl-CoA and Oxaloacetate (47). We did not observe significant changes in citrate synthase activity for UVA and UVB exposures compared to unexposed controls (Figure 1d), indicating that mitochondrial content remains stable upon UV exposure. To assess the overall function of mitochondria and changes in electron transport chain (ETC) activity, we performed the Seahorse assay (48), where we monitored mitochondrial respiration and glycolysis in viable cells based on their oxygen OCR and ECAR under basal, i.e. before the addition of chemical modulators of ETC activity, and maximal conditions, i.e. after the addition of the protonophore FCCP to collapse the inner mitochondrial membrane gradient (Figure 1e-1f). We found that both UVA and UVB induced a significant reduction in OCR compared to unexposed controls in both basal and maximal conditions upon irradiation exposure levels of 60-100 J/cm^2^ of UVA and 1-3 J/cm^2^ of UVB (Figure 1e). ECAR was also measured as a proxy for glycolytic activity by acidifying the media through lactate diffusion. We observed that UVA exposure levels of 60–100 J/cm^2^ and UVB levels of 1–3 J/cm^2^ significantly reduced glycolytic activity (Figure 1f). Overall, both UVA and UVB significantly perturbed cells, increasing levels of CD within the mitochondrial genome and leading to an overall decrease in cellular fitness.

### UVA, but not UVB, stimulates ROS that promotes mtDNA base oxidation and the CD

To characterize the relationship between UV-induced oxidative stress and CD formation, we monitored ROS and mtDNA base damage upon UVA and UVB exposure in human skin fibroblasts. Using the irradiation protocol shown in Figure 1a, we exposed cells to 60, 80, or 100 J/cm^2^ of UVA or 1, 2, or 3 J/cm^2^ of UVB and quantified cellular ROS levels using the fluorescent H_2_DCFDA (DCF) assay (49). We observed that UVA, but not UVB, promoted ROS accumulation (Figure 2a). The highest increase in ROS was a 10-fold increase detected at 60 J/cm^2^ UVA irradiation, which decreased with higher UVA irradiation levels (80 and 100 J/cm^2^). We measured mtDNA base oxidation and photoproducts following UVA and UVB exposure with a dot blot assay employing an anti-8-oxo-dG and an anti-CPDs specific antibody, respectively. We observed a 10-fold increase in 8-oxo-dG in fibroblasts irradiated with 100 J/cm^2^ of UVA as compared to unexposed cells (Figure 2b). Conversely, there was no detectable increase in CPDs upon UVA irradiation (Figure 2b). Regarding UVB-irradiated fibroblasts, we detected no monitorable change in 8-oxo-dG content, however, a 10-fold increase in CPD was detected (Figure 2b).

**Figure 2.**
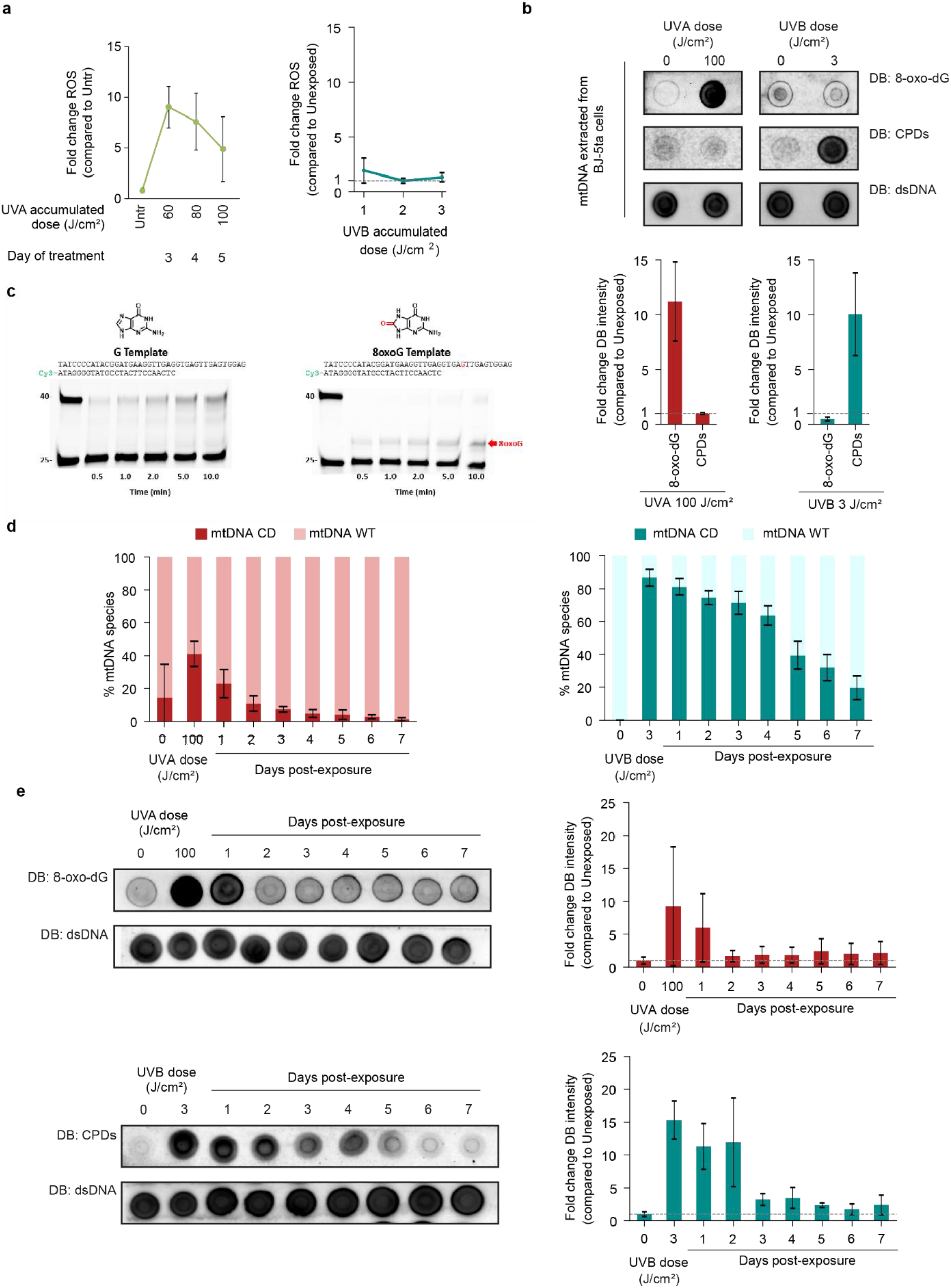
UV radiation induce distinct patterns of mtDNA base damage, leading to diverse kinetic profiles for the recovery of the mtDNA WT pool in absence of further irradiations. **(a)** Fold-change of cellular ROS content in BJ-5ta cells quantified by measuring the fluorescent 2’, 7’ –dichlorofluorescein (DCF) in UVA-(left) and UVB-exposed (right) cells to unexposed cells (grey dotted line). n = 3 biological replicates and analyzed by paired t-test comparing to unexposed control. **(b)** Representative dot blots (DB, top) and relative DBs quantitation (bottom) in fold change compared to unexposed cells (dotted grey line). DBs were performed on mtDNA extracted from BJ-5ta cells irradiated with the indicated doses of UVA (left) or UVB (right). Hybridization was performed with an anti-8-oxo-dG, an anti-cyclobutane pyrimidine dimer (CPDs), and an anti-double stranded DNA (dsDNA) antibody as a loading control. **(c)** Primer extension assays with mitochondrial DNA Pol γ replication past 40mer templates containing either an unmodified G Template (left) or a modified 8-oxo-dG template (right). Templates were pre-annealed opposite 25mer primers containing a 5’ Cy3 dye. **(d)** BJ-5ta cells were exposed to the indicated doses of UVA (left) and UVB (right) according to the scheme in Figure 1a. mtDNA samples were collected every 24 h for seven days after exposure in absence of further stimuli. Relative quantification of wild-type mtDNA (mtDNA WT, dark bars) and mtDNA harboring the common deletion (mtDNA CD, light bars) by qPCR in unexposed and UVA-(left) or UVB-exposed (right) cells. **(e)** Representative DBs (left) and relative DBs quantitation (right) in fold change compared to unexposed cells (dotted grey line). DBs were performed on mtDNA extracted from BJ-5ta cells irradiated with the indicated doses of UVA (top) or UVB (bottom) and collected every 24 h for seven days after exposure in absence of further stimuli. Hybridization was performed with an anti-8-oxo-dG, an anti-cyclobutane pyrimidine dimer (CPDs), and an anti-double stranded DNA (dsDNA) antibody as a loading control. n = 3 biological replicates for (b, d-e). Error bars represent standard deviations.

To further pursue the hypothesis that mtDNA base damage leads to replication fork stalling as an initiating event for the CD formation, we performed *in vitro* primer extension analysis to assess whether 8-oxo-dG leads to mtDNA replication stalling. We observed stalling of DNA polymerase Â on a 40-mer linear template at the 8-oxo-dG modification site (Figure 2c). In this time course study, maximal stalling was observed at the latest time point, ∼10 % stalling at 10 min. For the non-damaged template, full length synthesis was observed as function of increasing time, ∼15 % extension at 10 min. These results are in line with previously observed mitochondrial DNA polymerase Â stalling in the presence of 8-oxo-dG on both a 70-mer linear and minicircle template (50).

To assess whether UV-induced CD persists in the absence of further irradiation, we performed time course experiments monitoring CD levels up to seven days after UV exposures. The CD was present in the mtDNA pool (∼40 %) after an accumulated dose of 100 J/cm^2^ UVA. These levels steadily decreased after acute UVA stimulation, from ∼22 % on day 1 post-exposure to ∼1 % on day 7 post-exposure (Figure 2d, left). An accumulated dose of 3 J/cm^2^ of UVB resulted in CD levels of ∼85 %, as observed in Figure 1b. In absence of further UVB stimulation, CD levels decreased to ∼20 % on day 7 post-exposure (Figure 2d, right). Therefore, we concluded that CD levels from UVA and UVB follow distinct kinetic profiles, possibly indicating differences in the repair capacity of UV-induced damage. To correlate these diverse kinetic profiles to the potential persistence of mtDNA base damage in the post-exposure days, we performed dot blots monitoring base oxidation and the presence of photoproducts following the same time course scheme (Figure 2e and Figure S3). Upon recovery from the accumulated UVA dose (100 J/cm^2^), 8-oxo-dG levels decrease to the same levels as the unexposed control 2 days post-exposure (Figure 2e, upper panel). Similarly, the measured CPD levels from UVB exposure (3 J/cm^2^) recovered to the initial level 6 days post-treatment (Figure 2e, lower panel). Overall, we observed that UVA induced an increase in cellular ROS levels and an accumulation of 8-oxo-dG in normal fibroblasts. The CD pool was depleted with a similar kinetics to 8-oxo-dG, suggesting a link between the formation of CD within mitochondrial DNA and 8-oxo-dG.

### Antioxidants reduce UVA-induced ROS and protect against formation of the CD

To further test the hypothesis that UVA-dependent increase in ROS causes the CD to accumulate, we tested the capacity of antioxidants to reduce UVA-promoted formation of CD. Since UVB did not induce changes in ROS, we focused on UVA for the following experiments. We evaluated the influence of the antioxidants GSH (51) or CoQ10 (52) on cellular ROS and CD levels. ROS levels in fibroblasts at the start (S) and end (E) of each day’s exposure were determined (Figure 3a). Supplementing GSH or CoQ10 to human fibroblasts reduced endogenous cellular ROS levels (Figure 3b), although more prominently for GSH. The vehicles (H_2_O and DMF) did not have a significant effect on the endogenous cellular ROS levels, however, the presence of DMF delayed the induction of UVA-induced ROS (Figure 3b, right, day 2 end to day 3 start). The presence of GSH and CoQ10 reduced UVA-promoted ROS levels at similar levels at the end of 5 days, although the kinetics of both antioxidants were distinct.

**Figure 3.**
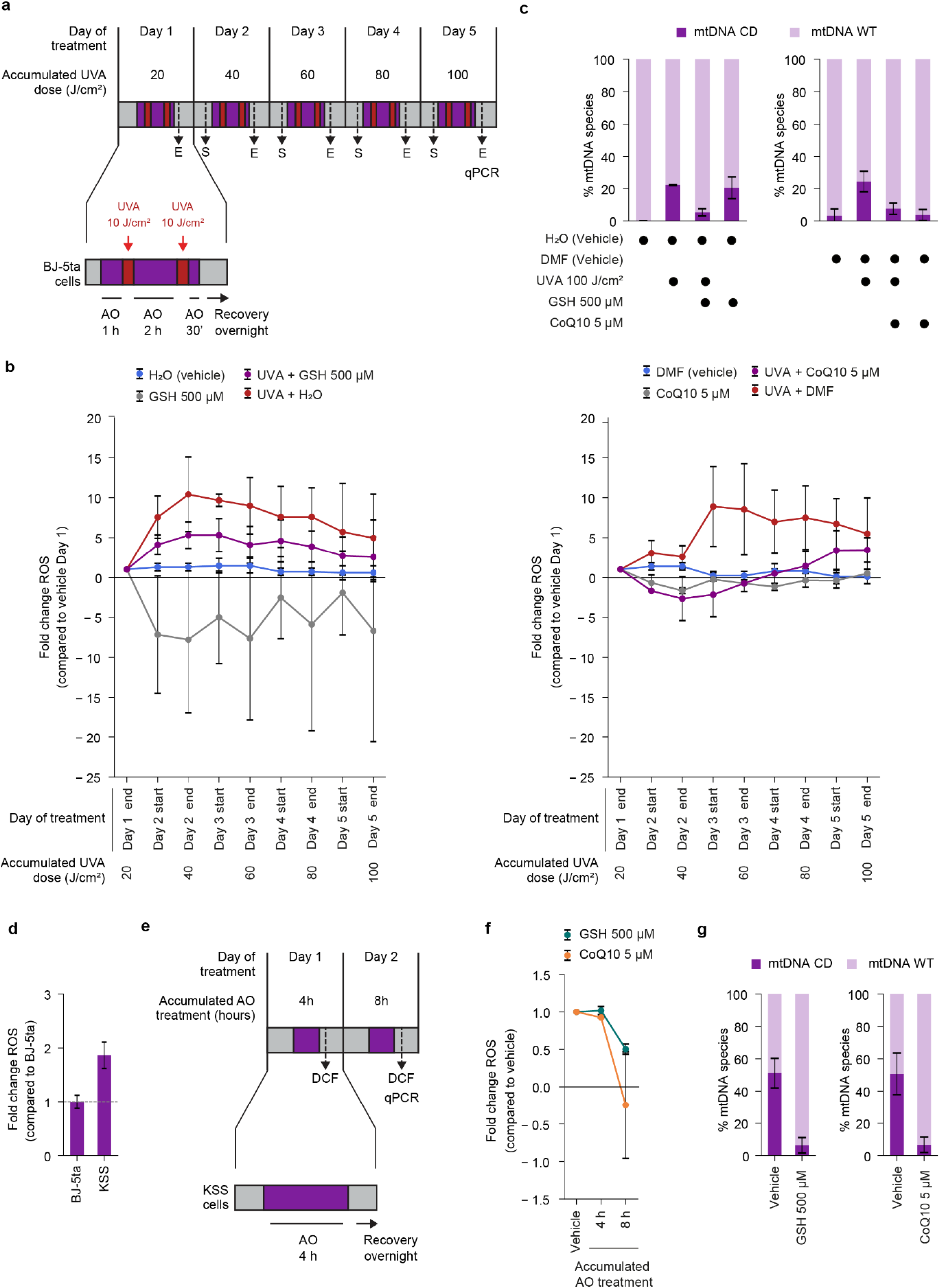
Antioxidant-mediated modulation of UVA-induced cellular ROS content reduces the formation or accumulation of the mtDNA CD. **(a)** Schematic representation of the UVA and antioxidants (AO) co-treatments performed on the BJ-5ta human fibroblasts cell line. Cells were pre-incubated with AO for 1 h in complete medium (purple bar). The AO-containing medium was removed, and cells were irradiated twice with UVA 10 J/cm^2^ in medium devoid of FBS and antibiotics (red bars). In between irradiations and after the last irradiation cells were incubated with AO for 2 h and 30 minutes, respectively. Harvesting (dotted arrows) was performed 30 minutes before and after the first last AO incubation, assessing the ROS content at the start (S) or end (E) of the exposure regime. Sampling for qPCR-based assessment of the mtDNA CD content is also indicated. **(b)** Cellular ROS content in BJ-5ta under-going the co-treatment with reduced glutathione (GSH, left), Coenzyme Q10 (CoQ10, right) and/or the corresponding vehicles H2O or Dimethylformamide (DMF). Cells were collected at the start and at the end of the UV exposure procedure and the ROS content was quantified using the DCF assay. Data are expressed as fold change to cells exposed to vehicle at the end of the first day. **(c)** Relative quantification of wild-type mtDNA (mtDNA WT, light bars) and mtDNA harboring the common deletion (mtDNA CD, dark bars) by qPCR of BJ-5ta cells exposed to UVA and co-treated with glutathione (GSH, left), Coenzyme Q10 (CoQ10, right) and/or the corresponding vehicles H_2_O or Dimethylformamide (DMF) as in the scheme in Figure 3a. Left panel: n = 3 and right panel n = 2. **(d)** Basal levels of ROS in BJ-5ta cells and human fibroblasts from a Kearns-Sayre syndrome (KSS) patient displaying inherent levels of oxidative stress. Fold change of cellular ROS content expressed as fold-change to ROS content in BJ-5ta cells (grey dotted line). **(e)** Schematic representation of the AO treatments in KSS cells. Cells were acutely exposed to AO for 4 h per day in complete medium, for 2 days. Cells were harvested 30 mins after the last incubation and their ROS and mtDNA CD contents were assessed by DCF and qPCR assays, respectively. **(f)** ROS content in KSS cells treated with vehicle or with GSH or CoQ10 for 4 h per day over 2 days. **(g)** Relative quantification of mtDNA WT (light bars) and mtDNA CD (dark bars) by qPCR in KSS cells treated with vehicle or with the indicated AO for 8 h, 4 h per day. n = 3 biological replicated for panel **(b-g)**. Error bars represent standard deviations.

Having established that both antioxidants have a similar ability to counteract UVA-promoted ROS, we measured CD levels by qPCR for the same conditions. UVA exposure at 100 J/cm^2^ in the presence of the vehicle increased CD levels to 22 % (Figure 3c, left), surprisingly also in the presence of GSH without UVA exposure. UVA-promoted CD formation was reduced to 5% by addition of GSH before and after UVA irradiation compared to UVA exposure and no GSH addition. Similarly, for the CoQ10 antioxidant, the addition of the antioxidant led to a reduction in the CD level from ∼22 % (UVA-exposed without CoQ10) to ∼5 % with the addition of CoQ10 (Figure 3c, right). These data illustrate that UVA-promoted ROS can directly be linked to the formation of the CD, which can be counteracted with the addition of antioxidants in human fibroblasts.

To further confirm the relationship between the formation of the CD and ROS, we tested the consistency of the previously stated findings in a more physiologically relevant model of human pathology. We evaluated changes in ROS and CD levels in a skin cell line derived from a KSS patient’s skin biopsy, which was enucleated and fused to a human rh0 osteosarcoma cell line (53), resulting in its capacity to stably maintain the pathological CD state. The basal ROS levels in the KSS cell line were 2-fold higher than in the BJ-5ta normal fibroblasts (Figure 3d). After the addition of the antioxidants GSH (500 µM) and CoQ10 (5 µM) during a 2-day transient exposure (Figure 3e), a robust reduction in ROS was observed at 8 h, especially for CoQ10 (Figure 3f). While in these KSS hybrid cells, a stable level of CD is present at ∼50 % heteroplasmatic content in their basal state, it was nearly eliminated by the addition of both GSH and CoQ10 over 8h (Figure 3g). Overall, these data support the ability of UVA-induced (Figure 3a-c) or endogenous (Figure 3e-g) oxidative stress to cause CD accumulation, which can be reduced by the presence of antioxidants.

### UVA exposure increases the expression of mtDNA replication genes and concomitantly decreases the levels of mtDNA-encoded genes

Having established that UVA induces CD through ROS production and DNA base oxidation, we aimed to explore the resulting changes in gene expression and associated pathways. Thus, we performed transcriptomic analysis of nuclear and mitochondrial genes in UVA-exposed fibroblasts on bulk RNA from 5 biological replicates of each condition. A stringent analysis of global gene expression changes evident from RNA-seq analysis revealed that among the 21,505 genes monitored, 6,372 genes were differentially expressed due to the UVA exposure (p<0.01k correlation of log ratio > 0.5 between biological replicates). Gene ontology analysis on the 2,000 most up- and down-regulated genes (Figure 4a) led to the identification of 6 gene clusters: 3 with up-regulated genes, and 3 with down-regulated genes. Interestingly, DNA damage and repair and modulation of the cell cycle were the most deregulated pathways. We then evaluated genes encoding known effectors involved in mtDNA maintenance, and in mtDNA replication, repair and degradation, revealing that genes involved in mtDNA replication are generally upregulated by UVA, while genes involved in mtDNA repair are mostly downregulated (Figure 4b). By monitoring genes relevant for the corresponding pathways in the nuclear genome, we found a similar downregulation for genes encoding for DNA repair (Figure S4a). However, genes encoding enzymes involved in nuclear DNA replication exhibited an opposite trend compared to those on the mitochondrial genome, which were also downregulated by UVA (Figure S4b). We used qPCR to validate the RNA-seq results for genes involved in mtDNA maintenance, e.g. mtDNA degradation, mtDNA replication and repair (base excision repair (BER), mismatch repair (MMR) and double-strand break repair (DSBR)) and found a good consistency between both methods, again showing that genes involved in mtDNA replication and degradation were generally upregulated upon UVA irradiation (100 J/cm^2^), whereas most repair-related genes were downregulated or unchanged (Figure 4c).

**Figure 4.**
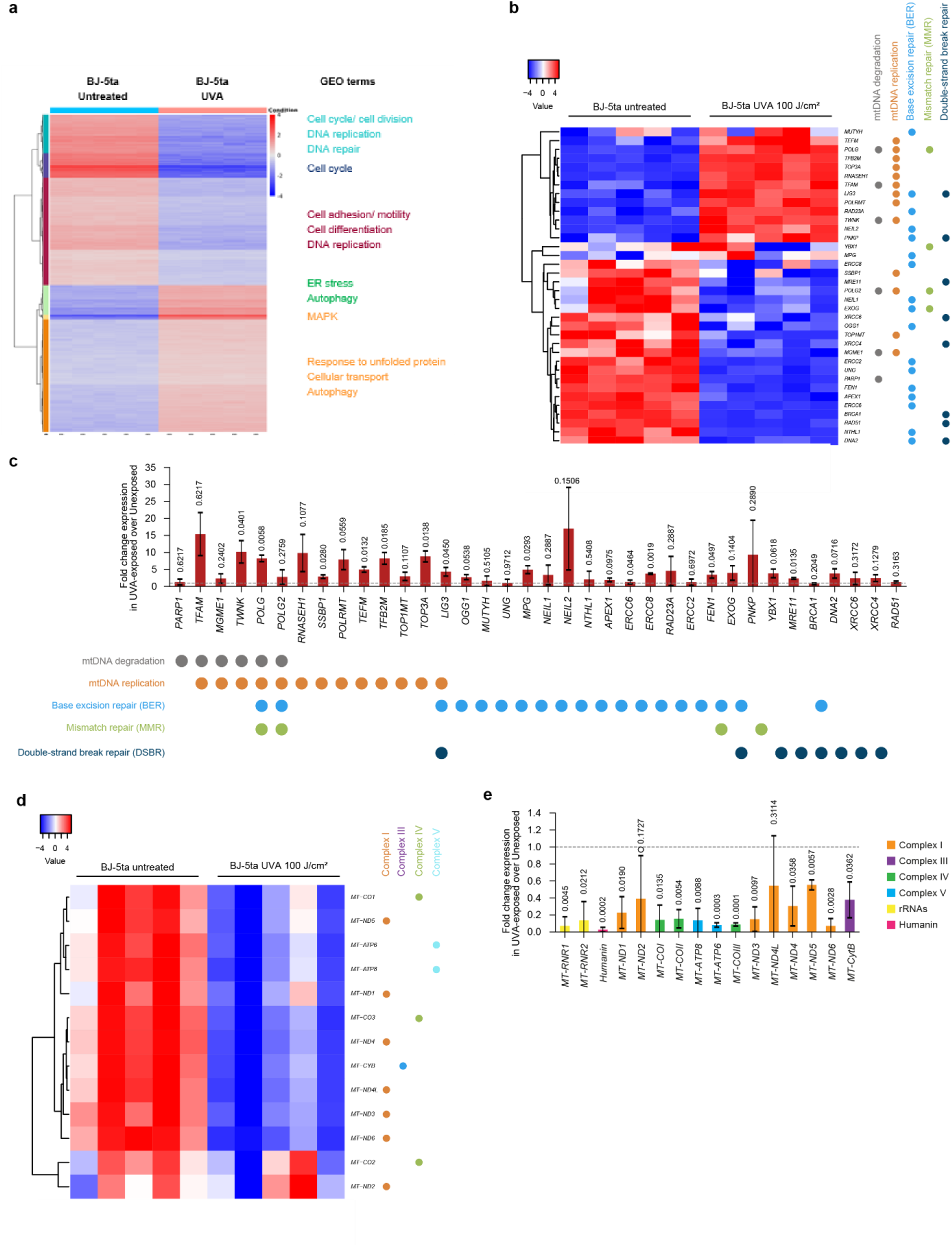
UV-irradiation induces gene expression changes in nuclear DNA-encoded mtDNA maintenance genes and in mtDNA-encoded genes. **(a)** RNA-seq results depicting the genes most affected in their expression upon exposure of BJ-5ta cells with UVA 100 J/cm^2^ according to the scheme in Figure 1a. 2000 most up- and downregulated genes were binned in 6 clusters (colored bars, left) and GEO terms were assigned to each cluster (right). **(b)** RNA-seq analysis of genes involved in mtDNA maintenance (mtDNA replication, repair and degradation) in unexposed and UVA-exposed BJ5-ta fibroblasts. **(c)** qPCR assays for expression of mtDNA maintenance genes upon UVA-exposed cells. n = 3 biological replicates were used for each graph. Statistical analysis was performed with a paired t-test comparing UV-exposed samples to matched unexposed control. **(d)** RNA-seq analysis of mtDNA-encoded genes in five unexposed and UVA-treated biological replicates. **(e)** qPCR analysis for the expression of mtDNA-encoded genes in UVA-exposed cells, expressed as fold change over unexposed cells (grey dotted line). Color codes indicate the ETC complexes to which the encoded proteins belong to, rRNAs and the Humanin micropeptide. n = 3 biological replicates were used for each graph. Statistical analysis was performed with a paired t-test comparing UV-exposed samples to matched unexposed control.

RNA-seq analysis further revealed that all 13 protein-coding mtDNA genes were downregulated by UVA irradiation (100 J/cm^2^, Figure 4d). The genes encoding the mitochondrial ribosomal RNA subunits (*MT-RNR1* and *MT-RNR2*, encoding the 12S and 16S rRNAs, respectively) were also downregulated by UVA. As a control, we evaluated gene expression changes in response to UVB irradiation (3 J/cm^2^) for the same genes shown in Figures 4c and 4e. While we observed similar deregulation in mtDNA-encoded genes, the expression changes in nuclear-encoded genes involved in mtDNA maintenance were specific to the type of UV exposure (Figure S5a-b). Overall, we observed that UVA exposure led to the upregulation of genes involved in mtDNA degradation, the downregulation of repair-related genes, and a reduction in mtDNA-encoded gene expression.

### UVA induces CD formation in keratinocytes and fibroblasts in a 3D human skin equivalent model

Having established in a 2D cell line that UVA promotes ROS generation, is associated with increased CD formation and is counteracted by antioxidants, we tested the potential of the CD as a biomarker of UVA-induced effects in a more physiologically relevant model. Therefore, we used a 3D tissue-engineered, organotypic full-thickness human skin equivalent (HSE) model. To construct the HSE, human primary keratinocytes were cultured on a natural dermis-like compartment. The dermis-like compartment is derived from triple seeding of human primary fibroblasts and their synthesis of an extracellular matrix within of four weeks. To induce keratinocyte, the HSEs were exposed to the air-liquid interface after three days. This model results in epidermal and dermal layers that are histologically similar to native human skin. Since this model is constructed using a fibroblast-derived extracellular matrix instead of plastic, it provides a more physiologically relevant microenvironment for the skin cells (44). The UVA exposure regime of the HSEs involved sequential irradiations (Figure 5a) where final exposure levels of 40, 60 and 80 J/cm^2^ were achieved. Subsequently, mtDNA was isolated and qPCR analysis was performed to quantify levels of the CD. Further, the CD levels were determined separately in both the epidermal and dermal layers, i.e. in keratinocytes and fibroblasts, respectively.

**Figure 5.**
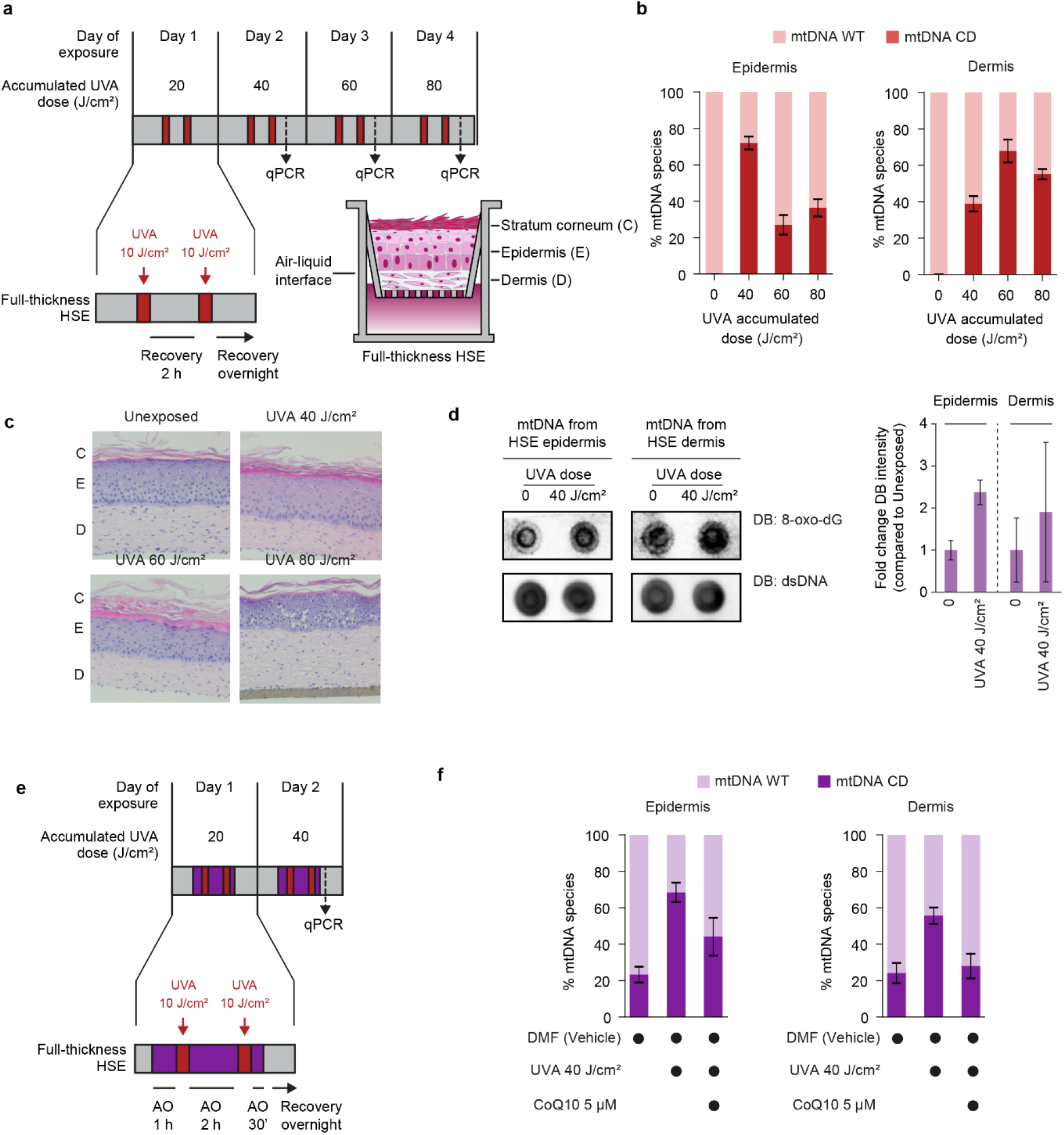
UVA irradiation induces ROS-dependent accumulation of the mtDNA CD in the epidermis and dermis of full-thickness HSE. **(a)** Schematic representations of the UVA exposure regime performed on three-dimensional full-thickness human skin equivalents (HSE). HSE were irradiated twice per day with the indicated doses of UVA (red bars), interrupted by recovery intervals (grey bars). Harvesting was performed 30 minutes after the last indicated irradiation. Full-thickness HSE (inlet) are composed of primary human fibroblasts and keratinocytes, constituting the dermis (D) and epidermis (E) respectively, seeded on a plastic air-lifted scaffold. Cellular proliferation and interactions between the two skin layers lead to formation of the corneum layer (C), reconstituting the structure and functionality of human skin. **(b)** Relative quantification of wild-type mtDNA (mtDNA WT, light bars) and mtDNA harboring the common deletion (mtDNA CD, dark bars) by qPCR in HSE exposed to increasing doses of UVA. Dermis (left) and Epidermis (left) were analyzed separately. **(c)** Representative microscopy images of hematoxylin-eosin-stained HSE sections, in unexposed conditions or exposed to the indicated doses of UVA. **(d)** Left: dot blots performed with mtDNA extracted from the epidermis and dermis of HSE, untreated or treated with 100 J/cm^2^ UVA and hybridized with anti-8oxodG and anti-dsDNA antibodies. Right: quantitation of n=3 biological replicates of dot blots. **(e)** Schematic representation of the UVA and CoQ10 co-treatments performed on HSE. Cells were pre-incubated for 1 h with 5 µM CoQ10 in complete medium (purple bar). The CoQ10-containing medium was removed, and HSE were irradiated twice with UVA 10 J/cm^2^. In between irradiation and after the last irradiation cells were incubated with CoQ10 for 30 minutes, respectively. Harvesting (dotted arrows) was performed 30 minutes after removing the CoQ10-containing medium. **(f)** Relative quantification of mtDNA WT (light bars) and mtDNA CD (dark bars) by qPCR of HSE exposed to UVA and co-treated 5 µM CoQ10 and/or the corresponding vehicle scheme in Figure 5f. Dermis (left) and Epidermis (left) were analyzed separately. n = 3 biological replicated for panel **(b-g)** Error bars represent standard deviations.

Upon dissociation of the epidermal and dermal layer in the HSE, an increase in the CD upon UVA-exposure was observed in both skin layers (Figure 5b). The highest CD levels (∼70 %) were observed for keratinocytes exposed to the lowest UVA dose (40 J/cm^2^), whereas the relative CD levels decreased with higher UVA doses (60 and 80 J/cm^2^). For fibroblasts, the highest relative CD levels were reached at the dose of 60 J/cm^2^, (Figure 5b). To assess whether these exposure levels affected tissue integrity, hematoxylin-eosin staining was performed. The structure of the HSE was not significantly perturbed by 40 or 60 J/cm^2^ UVA, while separation of the epidermis from the dermis started to be observed at 80 J/cm^2^ (Figure 5c) Consistent with the findings in the fibroblast monolayer (Figure 2c), dot blot analysis using an anti-8-oxo-dG antibody showed an increase in 8-oxo-dG in both layers of the HSE (Figure 5d). Overall, UVA-induced CD formation was observed in keratinocytes and fibroblasts along with an increase in base oxidation, although the extent of CD levels and kinetics were distinct between the two skin layers.

To confirm the anticipated protective effects of antioxidants on the formation of the CD (Figure 3), we exposed HSE for two days to 2x 10 J/cm^2^ UVA, reaching a maximum accumulated irradiation exposure of 40 J/cm^2^ UVA, either in the presence or absence of CoQ10 (Figure 5e). Whereas UVA exposure increased CD in keratinocytes to a level of 70 %, CoQ10 reduced it to 45 %. In fibroblasts, CD levels following UVA exposure were 55 %, and were also reduced (∼28 %) by the presence of CoQ10 (Figure 5f). In conclusion, we demonstrated that UVA-induced mtDNA CD can be measured in keratinocytes and fibroblasts in a 3D skin model and that these effects can be mitigated by supplementation with antioxidants. These findings support further investigation of mtDNA CD as a potential biomarker for UVA-induced mitochondrial damage, while highlighting the potential of the HSE model for in vitro testing.

## DISCUSSION

In this study, we characterized the effects of UV radiation in human skin fibroblasts and evaluated the mtDNA as a potential biomarker of UV-induced skin changes. We observed that increasing levels of UV radiation led to the accumulation of the mtDNA CD in human skin fibroblasts. UVA irradiation, but not UVB, stimulated ROS, which in turn promoted mtDNA base oxidation and CD formation. We demonstrated that antioxidants reduce UVA-induced ROS and CD levels, supporting CD formation is driven by UVA-induced ROS and could be counteracted by interventions that reduce ROS formation. Furthermore, UVA irradiation reduced the expression of mitochondrial encoded genes, while upregulating mtDNA replication genes upon UVA irradiation. Finally, using a 3D skin model physiologically similar to human skin, and were able to quantify how CD content changes upon UVA exposure, counteracted by antioxidants, supporting the CD as a biomarker to assess skin aging *in vitro*.

While it is reported that, *in vitro* exposure of fibroblasts and keratinocytes to UVA (31) and UVB (54) induces the CD, and free radicals may contribute, the distinct roles of UVA- or UVB in this process have yet to be fully defined. Thus, we exposed fibroblasts to increasing doses of UVA and UVB, and assessed mtDNA content and cellular fitness (Figure 1). The UV radiation doses were at the high end of environmental relevant exposure levels and were defined based on initial dose-range studies, and validated by detecting upregulation of *MMP-1* and downregulation of *COL1A1*, which aligns with the role of *MMP-1* in cleaving *COL1A1* upon UV exposure, contributing to collagen degradation and UV-induced skin damage and photoaging (55,56).

The relative mtDNA CD content increase in a dose-dependent manner, with UVB exhibiting a more pronounced effect (Figure 1b). It is established that UV irradiation reduces oxygen consumption and ATP production, and therefore has the potential to affect cell migration and division (57). A significant decrease in cell viability was observed, yet mitochondrial density remained unaltered (Figure 1c-e). The results of the functional assays demonstrated a notable decline in both the OCR and ECAR for the two UV exposures, evaluated at both basal and maximal ETC activity levels. The decline in OCR, as previously reported for UVA-exposed fibroblasts (58), suggests that UV irradiation may induce mitochondrial dysfunction. Notably, mitochondrial dysfunction is one of the hallmarks of aging (27) and UVA has been associated with photoaging (59).

To further characterize how UV radiation initiates the formation of the CD, we measured ROS and base damage (8-oxo-dG and CPDs) upon irradiation. We detected significantly increased cellular ROS levels upon UVA irradiation, peaking at 60 J/cm^2^ (Figure 2a), consistent with findings reported in the literature (60,61). Consistent with these results, we also detected elevated levels of 8-oxo-dG in the mtDNA of UVA-irradiated cells (100 J/cm^2^). Interestingly, cellular ROS levels decrease between 80 and 100 J/cm^2^ whereas the CD content increases in a dose-dependent manner. This may be attributed to reduced mitochondrial metabolism (Figure 1e-f), which might decrease cellular ROS levels in the cells. Additionally, higher UVA doses might allow sufficient time for the upregulation of antioxidant enzymes such as superoxide dismutase and catalase, which counteract ROS (62). The observed decrease in cellular ROS in UVA-exposed cells could also be a result of p53-mediated initiation of apoptosis in cells harboring higher levels of ROS (63).

In contrast to UVA, we did not observe an increase in cellular ROS upon UVB exposure in fibroblasts. However, consistent with the ROS measurements, we did not observe a significant increase in 8-oxo-dG levels in mtDNA. We measured the expected increase in CPD levels (10-fold, Figure 2b) in UVB-irradiated fibroblasts compared to unexposed cells (64). In this study, we found that although UVA and UVB both increase relative CD levels in fibroblasts (Figure 1b), the associated base damage and cellular ROS behavior are distinct. Thus, UVA triggered increased levels of ROS and base oxidation, whereas UVB induces CD without altering cellular ROS. What seems to be consistent for both UVA and UVB, is that irradiation increased mtDNA base damage (8-oxo-dG for UVA and CPD for UVB) along with increasing levels of the CD, and that the kinetics of the decrease of the CD and base damage post-exposure is simultaneous. Our data clearly highlights the association between base damage and the CD levels in mtDNA, and therefore, we propose the existence of two distinct initiation processes leading to the UV-dependent formation of the CD in human skin fibroblasts: (i) Stalling of polymerase Â at 8-oxo-dG (50) (Figure 2c) or CPDs (65), which could allow the 13bp repeats to mis-hybridize and lead to loop formation and subsequent deletion formation either through replication slippage (23,24) or (ii) the copy choice recombination pathway (29).

To link the UVA-dependent increase in ROS directly to CD accumulation, we tested if antioxidants (500 µM GSH and 10 µM CoQ10) reduce UVA-induced CD formation (Figure 3). Indeed, we observed lower levels of CD in UVA-exposed cells supplemented with antioxidants. Interestingly, GSH-exposed cells had lower ROS levels, but was not accompanied by a consequent reduction in CD levels. Fang Ji et. al. (66) similarly observed that GSH higher than 200 µM induced more CD even in unexposed cells. This observation was attributed to the potential of GSH at high concentrations to act as an electron donor, thereby contributing to ROS generation (66). In our experimental setup, however, GSH supplementation significantly lowered cellular ROS content relative to baseline levels, suggesting that the increase in CD observed in unexposed cells exposed to GSH may be explained in that GSH induces mitochondrial CD through reductive stress. Indeed, reductive stress is known to impair mitochondrial function (67) and could therefore lead to increased mitochondrial CD independent of ROS. This supports the idea that mitochondrial CD reflects mitochondrial quality and activity, suggesting it as an easily measurable indicator of mitochondrial health. Overall, our data demonstrate that exposing fibroblasts with GSH or CoQ10 reduces UVA-induced CD formation. Additional to previous studies that indicated the involvement of ROS in the formation of the CD (24), we measured cellular ROS content along with the CD to confirm the association between ROS and the CD are indeed ROS dependent. Since we observed that antioxidants could prevent the UVA-dependent increase in ROS and CD simultaneously, we could show that ROS is directly involved in the formation of the CD.

KSS is a rare disease characterized by mitochondrial dysfunction and known to be caused by the mitochondrial deletions, including the CD (68). We used a KSS donor derived cybrid cell line cell line and confirmed the association between ROS and the accumulation of the CD. We observed higher ROS levels in the KSS cell line at basal levels compared to the normal human fibroblast cell line BJ-5ta, which complements previous observations by Majora et al., where increased ROS and CD levels were detected in KSS-derived cell lines compared to normal human fibroblasts (69). By co-incubating the pathological cell line for 8 h with the antioxidants GSH or CoQ10, a decrease in steady-state ROS was detected. At the same time, we observed a decrease in CD content in steady-state CD levels in antioxidant co-treated cells. The mechanism describing how these high CD levels can persist in KSS is not yet known, however our data points to a connection between high content of ROS in basal state and the persistent accumulation of the CD in KSS cells. The mechanism for CD formation appears to be similar to UVA-induced CD formation, as CD formation can be reduced by antioxidant supplementation. This is consistent with a potential therapeutic role of antioxidant supplements for KSS patients, although the efficacy of remains to be established.

A key finding from the novel transcriptomic analysis presented here is the upregulation of genes associated with mtDNA replication and degradation, accompanied by the downregulation of genes linked to repair pathways (Figure 4). This observation suggests that, upon UVA irradiation, cells prioritize mtDNA turnover over repair. The upregulation of mtDNA replication genes could be an indication of a compensatory mechanism to replace damaged mtDNA, as previously proposed in studies investigating oxidative stress responses (70). The observed downregulation of DNA repair-associated genes could be an adaptive response to avoid an overload of repair intermediates, which carry inherent risks of increasing genome instability and indirectly disrupting genome function or causing mutations. We further showed a downregulation of all 13 protein-coding mtDNA genes upon UVA irradiation including the mitochondrial ribosomal RNA subunits (*MT-RNR1 and MT-RNR2*), which are essential for mitochondrial protein synthesis. The reduced expression of mitochondrial-encoded genes can be linked with compromised oxidative phosphorylation and enhanced ROS production, leading to a feedback loop of oxidative damage and impaired mitochondrial function. Our comparative analysis with UVB exposure revealed that while mtDNA-encoded genes showed similar patterns of downregulation, nuclear-encoded genes involved in mtDNA maintenance showed distinct expression profiles depending on the type of UV exposure. This suggests that UVA and UVB radiation trigger different transcriptional responses, likely due to differences in the type and extent of metabolic and DNA damage response.

In this study, we measured UVA-induced CD formation (Figure 5b) and mtDNA base damage (Figure 5d) in a physiologically relevant 3D skin model for the first time. Notably, we demonstrated that supplementing HSE with CoQ10 effectively mitigated the adverse effects of UVA-induced CD formation (Figure 5f), highlighting the potential of antioxidants as protective interventions against UV-induced damage and possibly photoaging. Furthermore, our data indicates the potential of HSE as an *in vitro* model for testing photoaging interventions, including antioxidant therapies. The observation that UVA-induced CD in keratinocytes (Figure 5b) decreases with increasing UVA exposure, together with upregulation of mtDNA replication genes upon UVA irradiation (Figure 4a) suggests there is a protective mechanism by which cells with high CD levels are replaced by active proliferation in keratinocytes. Thus, the data in this study supports that the UVA-specific initiating mechanism for elevated CD levels is by promotion of reactive oxygen species, supporting a molecular basis for how antioxidant interventions can mitigate UV-induced skin damage. Moreover, the 3D HSE model combined with CD monitoring technology establishes a basis for characterizing molecular and cellular responses to targeted interventions to prevent photoaging.

## DATA AVAILABILITY

All data generated during this study are available from the corresponding author upon reasonable request.

## SUPPLEMENTARY DATA

PDF contains Supplementary Figures 1–5, Supplementary Table 1.

## AUTHOR CONTRIBUTIONS

**G.A.F**. contribution: conceptualization, validation, methodology, data curation, formal analysis, supervision, visualization, writing original draft, review and editing. **N.K.S**. contribution: formal analysis, visualization, writing original draft, review and editing. **N.R**. contribution: methodology, data curation, formal analysis, visualization, writing original draft, review and editing. **A.E**-contribution: methodology, data curation, formal analysis, visualization, writing original draft, review and editing. **M.D.F**. contribution: methodology, data curation, formal analysis, visualization, writing original draft, review and editing. **M.M.A**. contribution: methodology, data curation, formal analysis, supervision, visualization, writing original draft, review and editing. **S.H**. contribution: conceptualization, review and editing. **F.W**. contribution: formal analysis, visualization, writing original draft, review and editing. **H.B.D**. contribution: conceptualization, writing original draft, review and editing. **S.J.S** contribution: conceptualization, writing original draft, review and editing. **H.J.G**. contribution: supervision, visualization, writing original draft, review and editing.

## ACKNOWLEDGEMENTS

We thank the ETH laboratory of toxicology for helpful discussion and feedback on this manuscript. Professor Ivan Tarassov from the University of Strasbourg for providing the KSS cell line. Genome-scale data produced and analyzed in this paper were generated using the resources of the Functional Genomics Center Zürich (FGCZ).

