## Supplementary File for "UVA irradiation promotes ROS-mediated formation of the common deletion in mitochondrial DNA"

***Table S1: Primers used in this study***

| Primers for qPCR-based analysis of mtDNA species | | | |
|---|---|---|---|
| **Target** | **Forward sequence (5'->3')** | **Reverse sequence (5'->3')** | **Reference** |
| *mtDNA total* | TAGCCCTAAACCTCAACAGT | TGCGCTTACTTTGTAGCCTTCAT | (1) |
| *mtDNA CD* | TTCCTCATCACCCAACTAAAAA | TTCGATGATGTGGTCTTTGG | (1) |
| *mtDNA undeleted* | TCGATGATGTGGTCTTTGGA | CATCTGTACCCACGCCTTCT | (2) |
| *ACTB genomic* | TCACCCACACTGTGCCCATCTACGA | CAGCGGAACCGCTCATTGCCAATGG | (2) |
| **Primers for qPCR-based analysis of gene level expression in cDNA** | | | |
| *ACTB genomic* | TCACCCACACTGTGCCCATCTACGA | CAGCGGAACCGCTCATTGCCAATGG | (2) |
| *APEX1* | GAGGAGCATGATCAGGAAGG | GCTGTTACCAGCACAAACGA | (2) |
| *BRCA1* | AGAGTCCAGCTGCTGCTCAT | CCCTGCTCACACTTTCTTCC | (2) |
| *DNA2* | AGCTCTTGGCATGAGTGAAAG | GCACGGTTAACTGTACAACAGC | (2) |
| *ERCC2* | TTCTCTGGGCTCGACGAC | AGTCGTACGGGAAGTAGACCAG | (2) |
| *ERCC6* | GGAGCAGAGGTGAAAATTGAA | CTCCTGGACAGGCATGAG | (2) |
| *ERCC8* | CAAGTCACAGACAAGAAATATTAGCAG | AGCACTTGCTGTTGCCAAG | (2) |
| *EXOG* | GGCTCCAGCAGGAAATAAC | CAAAATCCTGAGGCACAATG | (2) |
| *FEN1* | TCGAACTTGCTATGTAATTTGTGTC | AACTCAGCTGATTGCCAGGT | (2) |
| *GAPDH* | ACGGATTTGGTCGTATTGGG | TGATTTTGGAGGGATCTCG | (2) |
| *HUMANIN* | CGAGGGTTCAGCTGTCTCTT | GGCAGGTCAATTTCACTGGT | (2) |
| *LIG3* | CAACACGAAGACCCAGATCA | GGTACACATCACCGTGGAAA | (2) |
| *MGME1* | GACTGAAAAGCCCCAAAGTCT | GCGTGTCCTCCTTTCAGGTA | (2) |
| *MPG* | TTTACGGCATGTACTTCTGCAT | ATGGTCTCCAGACCTTCCAG | (2) |
| *MRE11* | ACAACCTGGAAGCTCAGTGG | TTAATACGCAGCAAACCAACA | (2) |
| *mtRNA UTR* | CTTTGATTCCTGCCTCATCC | TGATGTCTGTGTGGAAAGTGG | (2) |
| *MT-ATP6* | CCACAATCCTAGGCCTACCC | GGGATCAATAGAGGGGGAAA | (2) |
| *MT-ATP8* | GCCTACTCATTCAACCAATAGC | TCAGTAGAATTAGAATTGTGAAGATGA | (2) |
| *MT-CO1* | ATCCTACCAGGCTTCGGAAT | CGGAGGTGAAATATGCTCGT | (2) |
| *MT-CO2* | CCATCCCTACGCATCCTTTA | GGTCGCCTGGTTCTAGGAAT | (2) |
| *MT-CO3* | CCCGCTAAATCCCCTAGAAG | ATGGTGAAGGGAGACTCGAA | (2) |
| *MT-CYB* | TATCCGCCATCCCATACATT | GGTGATTCCTAGGGGGTTGT | (2) |
| *MT-ND1* | TGAAGTCACCCTAGCCATCA | GGTTCGGTTGGTCTCTGCTA | (2) |
| *MT-ND2* | AAGCAACCGCATCCATAATC | TCAGAAGTGAAAGGGGGCTA | (2) |
| *MT-ND3* | CCACAACTCAACGGCTACAT | TTGTAGGGCTCATGGTAGGG | (2) |
| *MT-ND4* | CTCGCTAACCTCGCCTTACC | AGTGAGCCCCATTGTGTTGT | (2) |
| *MT-ND4L* | TCGCTCACACCTCATATCCT | GCCATATGTGTTGGAGATTGA | (2) |
| *MT-ND5* | CGCTTCCCCACCCTTACT | GCGAGGGCTGTGAGTTTTAG | (2) |
| *MT-ND6* | TCTGAATTTTGGGGGAGGTT | CCACAGCACCAATCCTACCT | (2) |
| *MT-RNR1* | AAACGCTTAGCCTAGCCACA | CTTTACGCCGGCTTCTATTG | (2) |
| *MT-RNR2* | ACTTTGCAAGGAGAGCCAAA | TGGACAACCAGCTATCACCA | (2) |
| *MUTYH* | GTGTGTATCAGGGCCAACAG | GACTGCACGGAGAGGACAC | (2) |
| *OGG1* | TGTCACCTACCATGGCTTCC | AGGCCCAGCTTCCTGAGA | (2) |
| *NEIL1* | AGCTGCGCCTGATACTGAG | GCTGAAAAGAGCCGGACAT | (2) |
| *NEIL2* | GGGCAGCAGTAAGAAGCTACA | TGCAGGACCAACCTCACC | (2) |
| *NTHL1* | AGGTGCTGCTGTCACTGATG | TCTGCAGGATGCTGTCCA | (2) |
| *PARP1* | GGGATGACCAGCAGAAAGTC | CTGCCTTGCTACCAATTCC | (2) |

| | | | |
|---|---|---|---|
| *PNKP* | GACAGCATCTTTGTGGGAGAC | AAGGTTGAGGGCAAACAGG | (2) |
| *POLG* | CAGGTACCACCCTGGAGTC | CCCTGTTCGAGACAGTGCTT | (2) |
| *POLG2* | CACGAACTTTTACACATGTATCC | CAGAGAGAACACAAGGAACC | (2) |
| *POLRMT* | CATGTACAACGCCGTGATG | GGCATCCTTCACCATGAATAA | (2) |
| *RAD23A* | ATGCGGCAGGTGATTCAG | AGGGCCTTCAACTGTAAAAGC | (2) |
| *RAD51* | GGGAATTAGTGAAGCCAAAGC | TGGTGAAACCCATTGGAACT | (2) |
| *RNASEH1* | GGCCTTTGTCAGGAAATCTG | CTTTCGCCTCCGATTCTTGT | (2) |
| *SSBP1* | CATGAGTCCGAAACAACTACCA | ACAGGGTCCTGACCCACTC | (2) |
| *TEFM* | GAGAAAGCTCCTCAAACCAG | CAATTCTTCGAGTACCAAAAACG | (2) |
| *TFAM* | TGCAACTTCTGTGGAAGCAT | GAATCAGGAAGTTCCCTCCA | (2) |
| *TFB2M* | AAATTTGGACGAATAGAAGTAAATATG | AGTCTGGATTTCCGGGATCT | (2) |
| *TOP1MT* | CACAACAAAGGAGGTTTTCC | CCAGGCTCTTGATGACTTCC | (2) |
| *TOP3A* | GCCCAAGAGCAAGTGGCG | CCTCATGGTTTCTTTAGCATTT | (2) |
| *TWNK* | GGACCTGCCCCTCTATTTC | CATAGACGTAGACTGCATGTTGC | (2) |
| *UNG* | GCTGAGTGCCGAGCAGTT | TGCTTCTTCCAGCTCTCTCC | (2) |
| *XRCC4* | GAGGTAGGATCCGGAAGTGG | GAAACAAGGTGGATTCTGCTTA | (2) |
| *XRCC6* | GCCTTGTCCTCAGCCAGTTA | CCTCGACTTATGTCGGGTAGA | (2) |
| *YBX1* | CAATGTAAGGAACGGATATGG | GGTGTACAAATACATCTTCCTTGG | (2) |

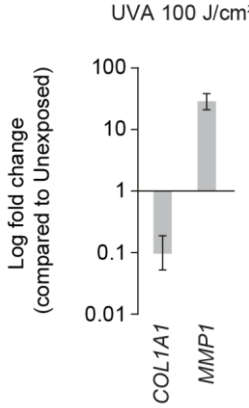


**Figure S1 | UVA-induced gene expression changes in UV responsive genes.** qPCR-based analysis of expression of *COL1A1 and MMP1* upon treatment of Bj5-ta cells with 100 J/cm$^2$ as shown Figure 1a. n = 3.

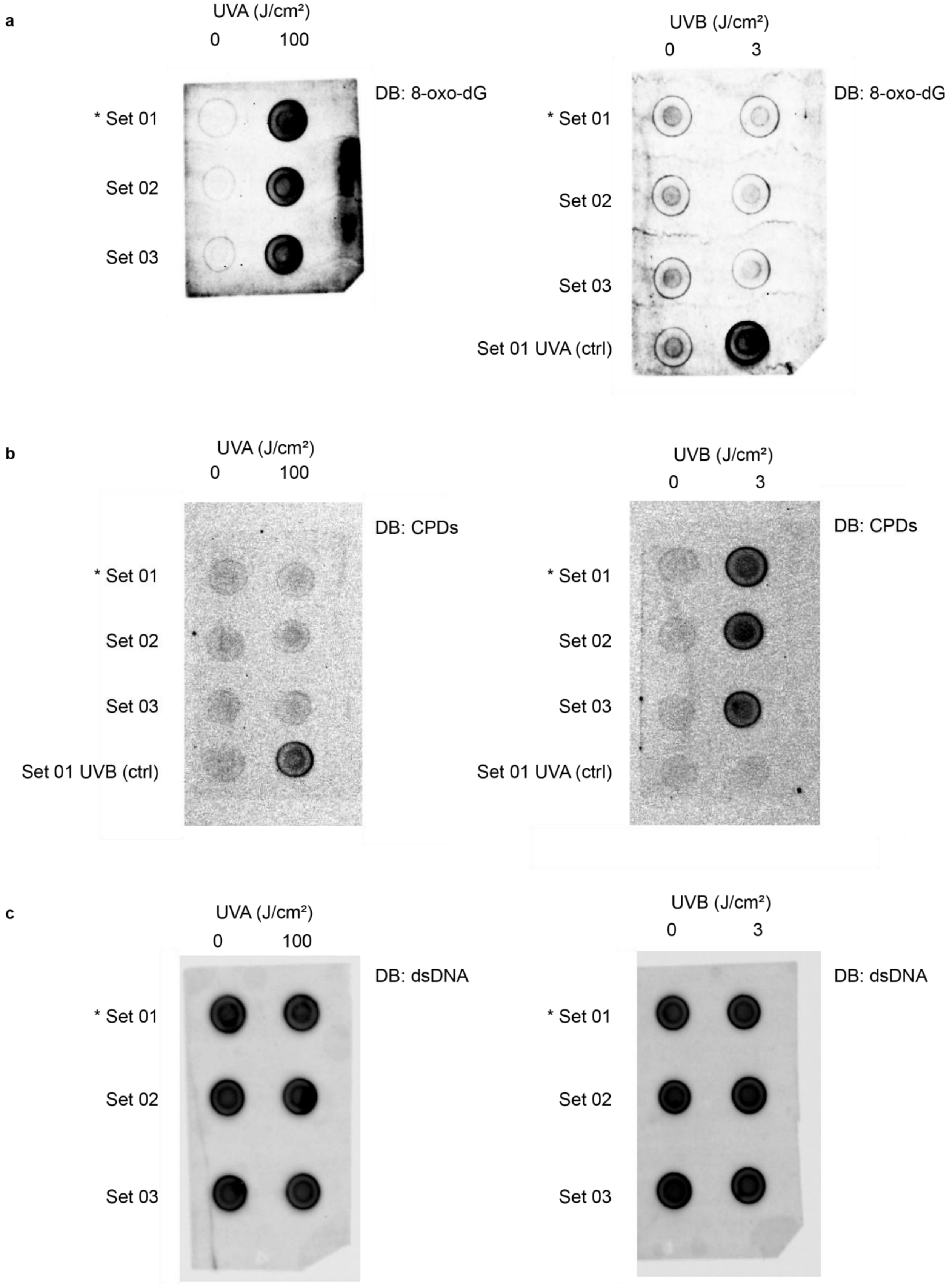


**Figure S2 | Uncropped dot blots.** Dot blot analysis was performed on mtDNA extracted from BJ-5ta cells irradiated with the indicated doses of UVA (left) and UVB (right). Hybridizations were carried out using **(a)** anti-8-oxo-dG, **(b)** anti-cyclobutane pyrimidine dimer (CPD), and **(c)** anti-double stranded DNA (dsDNA) antibodies as a loading control. Set 1, Set 2, and Set 3 represent three independent biological replicates.

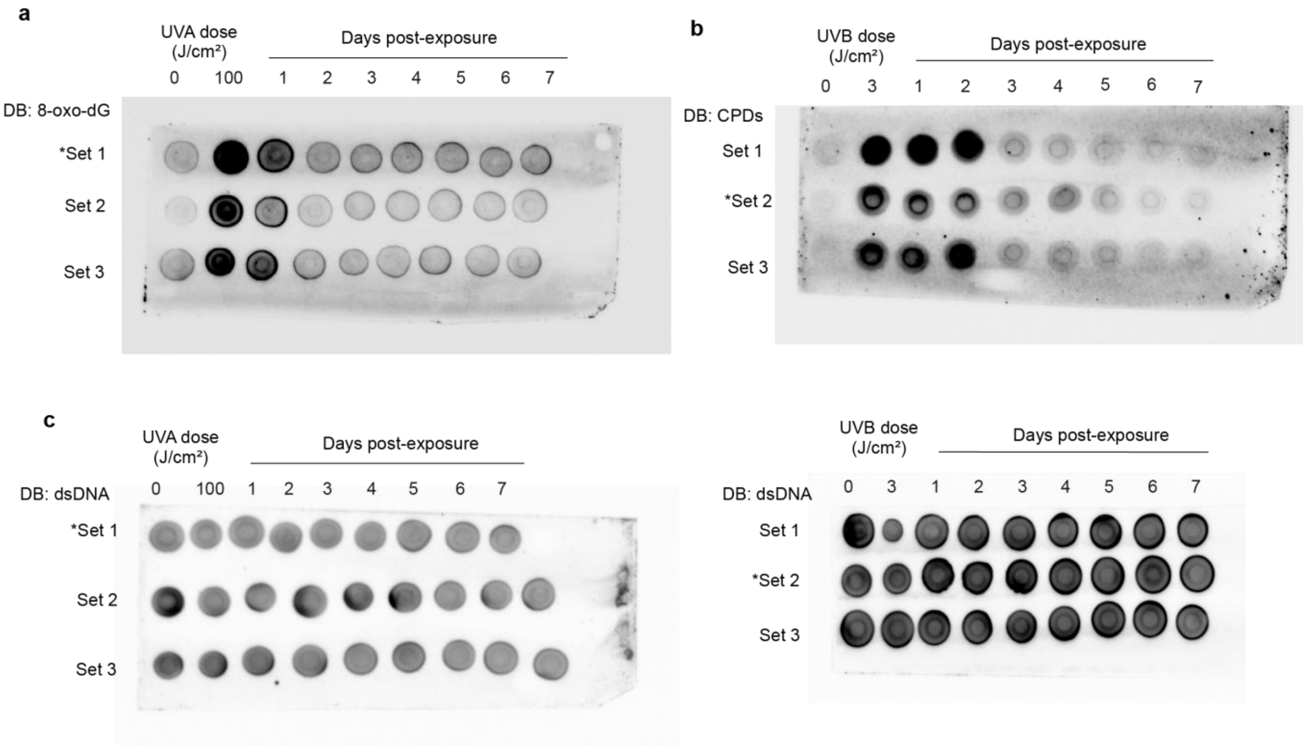


**Figure S3 | Uncropped dot blots.** DBs were performed on mtDNA extracted from BJ-5ta cells irradiated with the indicated doses of **(a)** UVA (100 J/cm$^2$) or **(b)** UVB (3 J/cm$^2$) and collected every 24 h for seven days after exposure in absence of further stimuli. Hybridizations were carried out using **(a)** anti-8-oxo-dG, **(b)** anti-cyclobutane pyrimidine dimer (CPD), and **(c)** anti-double stranded DNA (dsDNA) antibodies as a loading control. Set 1, Set 2, and Set 3 represent three independent biological replicates.

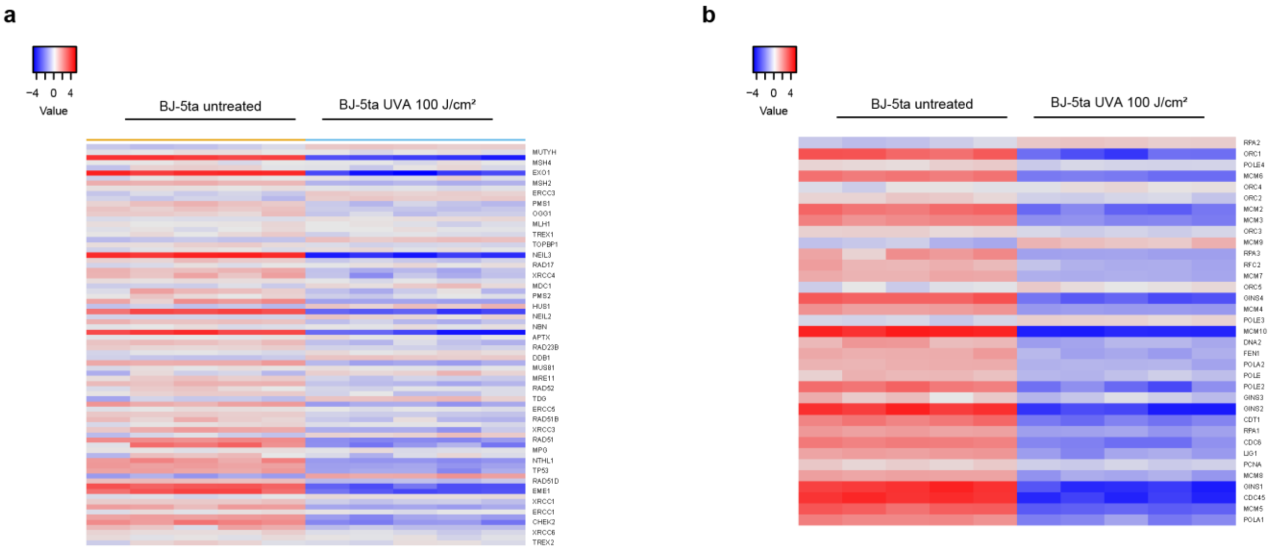


**Figure S4 | RNA-Seq analysis.** UV-irradiation (100 J/cm$^2$) induced gene expression changes in nuclear genes encoded for (a) DNA repair and (b) for DNA replication. n = 5

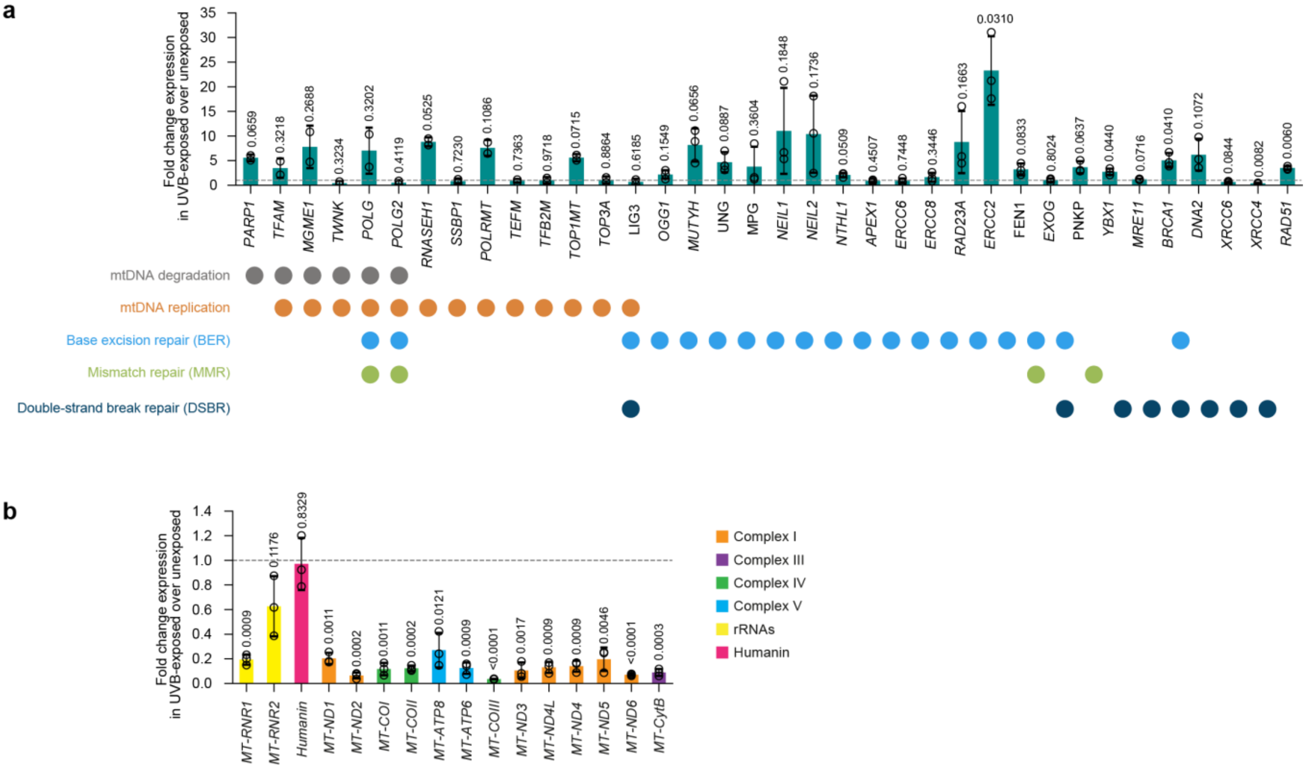


**Figure S5 | UVB-induced gene expression changes.** qPCR analysis of (a) nuclear-encoded mtDNA maintenance genes and (b) mtDNA-encoded genes in UVB-exposed cells. Data are expressed as fold change relative to unexposed cells (indicated by the grey dotted line). Statistical significance was determined using a paired t-test, comparing UVB-exposed samples to matched unexposed controls.

1. Phillips AF, Millet AR, Tigano M, Dubois SM, Crimmins H, Babin L, et al. Single-Molecule Analysis of mtDNA Replication Uncovers the Basis of the Common Deletion. Mol Cell. 2017;65(3):527-38.e6.

2. Fontana GA, MacArthur MR, Rotankova N, Di Filippo M, Beer HD, Gahlon HL. The mitochondrial DNA common deletion as a potential biomarker of cancer-associated fibroblasts from skin basal and squamous cell carcinomas. Sci Rep. 2024;14(1):553.
